# No evidence for associations between men’s salivary testosterone and responses on the Intrasexual Competitiveness Scale

**DOI:** 10.1101/198424

**Authors:** Jaimie S Torrance, Amanda C Hahn, Michal Kandrik, Lisa M DeBruine, Benedict C Jones

**Affiliations:** Institute of Neuroscience & Psychology, University of Glasgow, UK.; Department of Psychology, Humboldt State University, USA.; Institute for Brain and Behavior Amsterdam, VU Amsterdam, NL.

## Abstract

Many previous studies have investigated relationships between men’s competitiveness and testosterone. For example, the extent of changes in men’s testosterone levels following a competitive task predicts the likelihood of them choosing to compete again. Recent work investigating whether individual differences in men’s testosterone levels predict individual differences in their competitiveness have produced mixed results. Consequently, we investigated whether men’s (N=59) scores on the Intrasexual Competitiveness Scale were related to either within-subject changes or between-subject differences in men’s testosterone levels. Men’s responses on the Intrasexual Competitiveness Scale did not appear to track within-subject changes in testosterone. By contrast with one recent study, men’s Intrasexual Competitiveness Scale also did not appear to be related to individual differences in testosterone. Thus, our results present no evidence for associations between men’s testosterone and their responses on the Intrasexual Competitiveness Scale.

Data files and analysis scripts are publicly available at https://osf.io/abqun/

## Introduction

Results of several studies suggest that increases in men’s testosterone levels due to competitive tasks are associated with increases in their intrasexual competitiveness (reviewed in Zilioli & Bird, 2017). For example, men whose testosterone levels increased after competing against another man on a laboratory task (the Point Subtraction Aggression Paradigm, see Geniole et al., 2017 for a review of this method) were more likely to choose to compete again than were men whose testosterone levels did not increase after competing on the initial task (Carré & McCormick, 2008). Similarly, the extent to which men’s testosterone increases after losing a competitive task against another man is positively related to their willingness to compete again (Carré et al., 2009; Mehta & Josephs, 2006). These effects can be modulated by the decisiveness of the victory (Mehta et al., 2015a) and/or men’s aggressiveness (Carré & McCormick, 2008).

More recently, it has been hypothesized that some associations between testosterone and competition-related behaviors are moderated by cortisol (see Mehta & Prasad, 2015, for a discussion of evidence for this "Dual Hormone Hypothesis”). For example, Mehta et al. (2015b) found that behavior in a competitive bargaining game was predicted by the interaction between changes in testosterone and cortisol in a sample of men and women. When cortisol decreased, testosterone increases led to greater earnings (Mehta et al., 2015b). By contrast, when cortisol increased, testosterone increases led to poorer earnings (Mehta et al., 2015b). Given these results, failure to consider the moderating role of cortisol could explain null and negative results for relationships between testosterone and competition-related behaviors in some studies (Mehta & Prasad, 2015).

The studies described above investigated effects of competition-induced changes in testosterone on competitiveness. Other studies have investigated putative correlations between individual differences in men’s competitiveness and testosterone. Results from these studies have been mixed, however. Apicella et al. (2011) found no evidence that men with higher testosterone levels showed greater competitiveness (measured by rate of self-selection into a competitive setting). Arnocky et al. (2018) recently reported that men with higher testosterone scored higher on Buunk and Fisher’s (2009) Intrasexual Competitiveness Scale. Buunk and Fisher (2009) defined intrasexual competitiveness as viewing “confrontations with same-sex individuals, especially in the context of contact with the opposite-sex, in competitive terms”.

The aim of the current study was to investigate whether within-subject changes in reported intrasexual competitiveness tracked within-subject changes in men’s testosterone levels. Since we collected these data, Arnocky et al. (2018) published their article. Consequently, we also used our data to test whether men reporting greater intrasexual competitiveness would have higher testosterone levels. Like Arnocky et al. (2018), we assessed intrasexual competitiveness using Buunk and Fisher’s (2009) Intrasexual Competitiveness Scale.

## Methods

### Participants

Fifty-nine heterosexual men participated in the study (mean age=22.06 years, SD=3.24 years). None of these men were currently taking any form of hormonal supplement or had taken any form of hormonal supplement in the 90 days prior to participation. Participants took part in the study as part of a larger project investigating hormonal correlates of voice and face perception (Kandrik et al., 2016, 2017).

### Procedure

Participants completed up to five weekly test sessions, which took place between 2pm and 5pm to minimize diurnal variation in hormone levels (Papacosta & Nassis, 2011). Fifty-five of the participants completed more than one test session, with forty-seven of the participants completing all five test sessions.

During each test session, participants provided a saliva sample via the passive drool method (Papacosta & Nassis, 2011). Participants were instructed to avoid consuming alcohol and coffee in the 12 hours prior to participation and to avoid eating, smoking, drinking, chewing gum, or brushing their teeth in the 60 minutes prior to participation.

In each test session, participants also completed Buunk and Fisher’s (2009) Intrasexual Competitiveness Scale (M=2.95, SD=0.98; reliability: Cronbach’s alpha=0.86). The Intrasexual Competitiveness Scale is a 12-item questionnaire on which participants indicate how applicable each item is to them using a one (not at all applicable) to seven (completely applicable) scale. Example items include, "I want to be just a little better than other men” and "I tend to look for negative characteristics in men who are very successful”. Scores on individual items are averaged to produce an overall score. Higher scores on this scale indicate greater intrasexual competitiveness. The order in which participants provided saliva samples and completed the questionnaire was fully randomized. Like Arnocky et al. (2018), these data were collected as part of a larger project. The project was approved by University of Glasgow’s Psychology Ethics Committee.

### Assays

Saliva samples were immediately frozen and stored at -32°C until being shipped, on dry ice, to the Salimetrics Lab (Suffolk, UK) for analysis. There they were assayed using the Salivary Testosterone Enzyme Immunoassay Kit 1-2402 (M = 177.5 pg/mL, SD = 42.2 pg/mL, sensitivity <1.0 pg/mL, intraassay CV=4.60%, inter-assay CV=9.83%) and Salivary Cortisol Enzyme Immunoassay Kit 1-3002 (M = 0.19 μg/dL, SD = 0.11 μg/dL, sensitivity<0.003 μg/dL, intra-assay CV=3.50%, inter-assay CV=5.08%).

Hormone levels more than three standard deviations from the sample mean for that hormone or where Salimetrics indicated levels were outside the sensitivity range of the relevant ELISA were excluded from the dataset (<1% of hormone measures were excluded for these reasons; one cortisol value and four testosterone values). The descriptive statistics given above do not include these excluded values.

For *current* hormone levels, values for each hormone were centered on their subject-specific means to isolate effects of within-subject changes in hormones. They were then scaled so the majority of the distribution for each hormone varied from -.5 to .5 to facilitate calculations in the linear mixed models. To calculate *average* hormone levels, the average value for each hormone across test sessions was calculated for each man. These values were then centered on their grand means and scaled so the majority of the distribution for each hormone varied from -.5 to .5. Plots of these values are given in our Supplemental Materials and show no evidence of skew.

### Analyses

We used a linear mixed model to test for possible effects of hormone levels on reported intrasexual competitiveness. Analyses were conducted using R version 3.3.2 (R Core Team, 2016), with lme4 version 1.1-13 (Bates et al., 2014) and lmerTest version 2.0-33 (Kuznetsova et al., 2013). Data files and analysis scripts are publicly available at https://osf.io/abqun/.

## Results

The dependent variable was Intrasexual Competitiveness Scale score. Predictors were current testosterone, current cortisol, and their interaction, and average testosterone, average cortisol, and their interaction. No covariates were included in the model. Random slopes were specified maximally following Barr et al. (2013) and Barr (2013). Full model specifications and full results for each analysis are given in our Supplemental Information. Results are summarized in Table 1. There were no significant effects.

**Table 1.** Summary of results for men’s hormone levels and reported intrasexual competitiveness.

|  | Estimate | Std. Error | df | t | p |
| --- | --- | --- | --- | --- | --- |
| **Current Testosterone** | 0.053 | 0.227 | 38.280 | 0.235 | 0.815 |
| **Current Cortisol** | 0.094 | 0.216 | 34.620 | 0.434 | 0.667 |
| **Current Testosterone x Current Cortisol** | 2.565 | 1.566 | 162.41 | 1.637 | 0.103 |
| **Average Testosterone** | 0.389 | 0.647 | 60.290 | 0.601 | 0.550 |
| **Average Cortisol** | -0.221 | 0.865 | 61.430 | -0.256 | 0.799 |
| **Average Testosterone x Average Cortisol** | -4.421 | 3.029 | 59.540 | -1.460 | 0.150 |

We also collected and analyzed anxiety questionnaire data. These analyses are reported in our Supplemental Materials and show that men reported greater anxiety in test sessions where their cortisol levels were high.

## Discussion

Our analysis of men’s reported intrasexual competitiveness revealed no significant relationships between reported intrasexual competitiveness and men’s hormone levels. We found no evidence that within-subject changes in men’s reported intrasexual competitiveness tracked changes in men’s current testosterone, current cortisol, or their interaction. We also found no evidence that between-subject differences in reported intrasexual competitiveness were related to men’s average testosterone, average cortisol, or their interaction. These latter null results are noteworthy because they do not replicate Arnocky et al’s (2018) recent finding of a positive correlations between reported intrasexual competitiveness and testosterone level.

There are several limitations to our study that should be acknowledged. First, the order in which participants completed the questionnaires and provided saliva samples was randomized, rather than being controlled. This was also the case in Arnocky et al. (2018). Reducing noise associated with the order in which participants completed these two components of the study might yet reveal links between hormone levels and men’s reported intrasexual competitiveness not apparent in the current study. Second, although previous studies have detected within-subject changes in reported intrasexual competitiveness using the Intrasexual Competitiveness Scale (Buunk & Massar, 2012; Cobey et al., 2013; Hahn et al., 2016), it is possible that it is better suited to detecting hormone-linked individual differences than it is to detecting hormone-linked within-individual differences. Further work investigating changes in competitiveness using other methods may yet reveal hormone-linked changes not apparent in the current study. Third, although we do not replicate Arnocky et al’s (2018) results for individual differences in testosterone and intrasexual competitiveness, they had a larger sample that we did (92 men vs 59 men).

Previous research has suggested that women’s intrasexual competitiveness increases when their testosterone levels are high (Cobey et al., 2013; Hahn et al., 2016). By contrast, we found no evidence that intrasexual competitiveness tracked changes in men’s testosterone levels. These two studies (Cobey et al., 2013; Hahn et al., 2016) used the same Intrasexual Competiveness Scale as our current study. Further work is needed to establish whether the differences in these results reflect a sex difference in the effects of testosterone on intrasexual competitiveness, a false negative in the current study, or false positives in the studies of women’s intrasexual competitiveness. Nonetheless, our null results provide little support for the Challenge Hypothesis of testosterone and competition in men.

In conclusion, we found no evidence that men with higher testosterone levels scored higher on the Intrasexual Competiveness Scale. Moreover, because we also found no evidence that within-subject changes in scores on this measure tracked changes in testosterone, it is unlikely that the null result is due to testosterone-linked within-subject changes in responses obscuring the between-subject relationship. Of course, these results may not necessarily generalize to other measures of competition in men, which may be related to testosterone in other ways. Further work using a wider range of competition measures would clarify this issue.

