## Supplementary Materials for "No evidence for associations between men’s salivary testosterone and responses on the Intrasexual Competitiveness Scale" (from another version of this preprint: version 2, 2018-03-29)

```
library(tidyverse)
```

```
## Warning: package 'tidyverse' was built under R version 3.3.2

## Warning: package 'tibble' was built under R version 3.3.2

## Warning: package 'dplyr' was built under R version 3.3.2
```

```
library(lmerTest)
library(psych)
sessionInfo()
```

```
## R version 3.3.1 (2016-06-21)
## Platform: x86_64-apple-darwin13.4.0 (64-bit)
## Running under: OS X 10.11.6 (El Capitan)
##
## locale:
## [1] en_GB.UTF-8/en_GB.UTF-8/en_GB.UTF-8/C/en_GB.UTF-8/en_GB.UTF-8
##
## attached base packages:
## [1] stats     graphics  grDevices utils     datasets  methods   base
##
## other attached packages:
##  [1] psych_1.7.8     lmerTest_2.0-32 lme4_1.1-12     Matrix_1.2-6
##  [5] dplyr_0.7.3     purrr_0.2.2     readr_1.0.0     tidyr_0.6.0
##  [9] tibble_1.3.4    ggplot2_2.1.0   tidyverse_1.1.1
##
## loaded via a namespace (and not attached):
##  [1] reshape2_1.4.2      splines_3.3.1       haven_1.0.0
##  [4] lattice_0.20-33     colorspace_1.2-7    htmltools_0.3.5
##  [7] yaml_2.1.13         chron_2.3-47        survival_2.40-1
## [10] rlang_0.1.2         nloptr_1.0.4        foreign_0.8-66
## [13] glue_1.1.1          RColorBrewer_1.1-2  modelr_0.1.0
## [16] readxl_0.1.1        bindrcpp_0.2        bindr_0.1
## [19] plyr_1.8.4          stringr_1.3.0       munsell_0.4.3
## [22] gtable_0.2.0        rvest_0.3.2         evaluate_0.10
## [25] latticeExtra_0.6-28 knitr_1.20          forcats_0.2.0
## [28] parallel_3.3.1      htmlTable_1.7       broom_0.4.2
## [31] Rcpp_0.12.12        acepack_1.4.1       scales_0.4.0
```

```
## [34] backports_1.1.2     Hmisc_4.0-0         jsonlite_1.1
## [37] gridExtra_2.2.1     mnormt_1.5-5        hms_0.2
## [40] digest_0.6.10       stringi_1.1.7       grid_3.3.1
## [43] rprojroot_1.3-2     tools_3.3.1         magrittr_1.5
## [46] cluster_2.0.4       Formula_1.2-1       pkgconfig_2.0.1
## [49] MASS_7.3-45         data.table_1.9.6    xml2_1.1.1
## [52] lubridate_1.6.0     assertthat_0.2.0    minqa_1.2.4
## [55] rmarkdown_1.9       httr_1.2.1          rpart_4.1-10
## [58] R6_2.2.2            nnet_7.3-12         nlme_3.1-128
```

### Load Data

```
data_hormones <- read_csv("hm_intrasexual_comp_anon.csv")
```

### Descriptives

#### The number of sessions completed per man

```
data_hormones %>%
  group_by(hm_id) %>%
  summarise(
    sessions = n_distinct(date)
  ) %>%
  group_by(sessions) %>%
  summarise(
    n = n()
  )
```

```
## # A tibble: 5 x 2
##   sessions     n
##      <int> <int>
## 1        1     4
## 2        2     2
## 3        3     2
## 4        4     4
## 5        5    47
```

#### Mean age for the sample

```
data_hormones %>%
  group_by(hm_id, date, age) %>%
  summarise(n = n()) %>%
  ungroup() %>%
  group_by() %>%
  summarise(
      mean_age = mean(age, na.rm = TRUE),
      sd_age =   sd(age, na.rm = TRUE),
      se_age =   se(age, na.rm = TRUE)
  ) %>%
  mutate_all(round, 4)
```

```
## Warning: package 'bindrcpp' was built under R version 3.3.2
```

```
## # A tibble: 1 x 3
##   mean_age sd_age se_age
##      <dbl>  <dbl>  <dbl>
## 1  21.9679 3.1446 0.1932
```

### Data Processing

#### Exclude hormone outliers

```
# calculate means and SDs
test_mean <- mean(data_hormones$test)
test_sd <- sd(data_hormones$test)
cort_mean <- mean(data_hormones$cort)
cort_sd <- sd(data_hormones$cort)

# set values > 3SD from the mean to NA
data_final <- data_hormones %>%
  mutate (
    test = ifelse (test > test_mean + 3*test_sd |
                   test < test_mean - 3*test_sd, NA, test),
    cort = ifelse (cort > cort_mean + 3*cort_sd |
                   cort < cort_mean - 3*cort_sd, NA, cort)
  )

# determine how many values were excluded
data_final %>%
  group_by(hm_id, date) %>%
  summarise(
    t = is.na(mean(test)),
    c = is.na(mean(cort))
  ) %>%
  ungroup() %>%
  select(t:c) %>%
  gather('hormone','na', t:c) %>%
  group_by(hormone) %>%
  summarise(
    'valid' = n() - sum(na),
    'excluded' = sum(na)
  ) %>%
  arrange(hormone)
```

```
## # A tibble: 2 x 3
##   hormone valid excluded
##     <chr> <int>    <int>
## 1       c   264        1
## 2       t   261        4
```

#### Calculate average hormones for each participant

```r
data_avg <- data_final %>%
  group_by(hm_id) %>%
  summarise(
    avg_test = mean(test, na.rm = TRUE),
    avg_cort = mean(cort, na.rm = TRUE)
  ) %>%
  group_by() %>%
  # divide by constants to make range approx -.5 to +.5 for lmer
  mutate(
    avg_test.s = (avg_test - mean(avg_test, na.rm=TRUE)) / 180,
    avg_cort.s = (avg_cort-mean(avg_cort, na.rm=TRUE)) / 0.5
  )
```

#### Centre and scale hormones

Centre hormones on subject-specific means, and bring values between -0.5 and 0.5

```r
# centre hormones within-subject
# divide by same constants above to make range approx -.5 to +.5 for lmer
data_scaled <- data_final %>%
  group_by(hm_id) %>%
  mutate(
    test.s = (test-mean(test, na.rm=TRUE)) / 180,
    cort.s = (cort-mean(cort, na.rm=TRUE)) / 0.5
  ) %>%
  ungroup() %>%
  left_join(data_avg, by="hm_id")
```

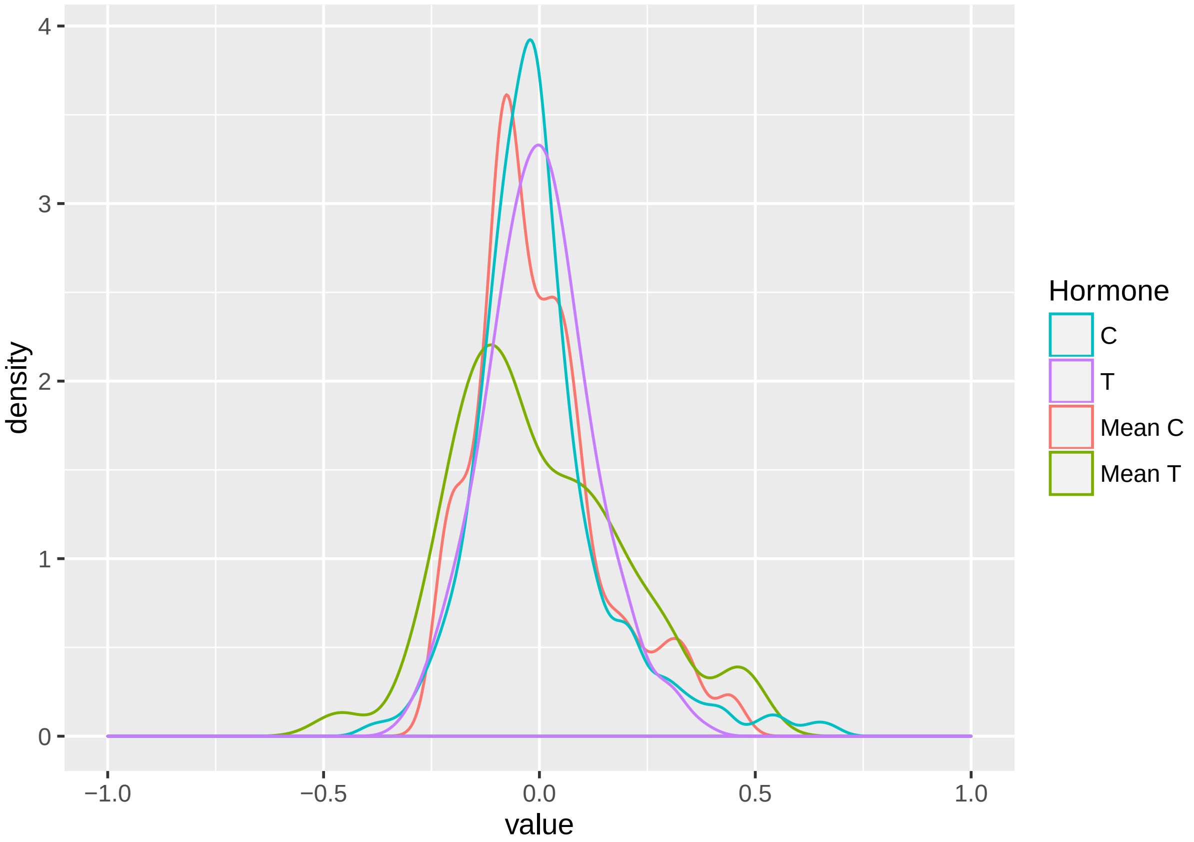

**Mean hormone levels**

```
data_scaled %>%
  group_by(hm_id, date, age, test, cort) %>%
  summarise(n = n()) %>%
  ungroup() %>%
  group_by() %>%
  summarise(
      mean_test = mean(test, na.rm = TRUE),
      sd_test =   sd(test, na.rm = TRUE),
      se_test =   se(test, na.rm = TRUE),
      mean_cort = mean(cort, na.rm = TRUE),
      sd_cort =   sd(cort, na.rm = TRUE),
      se_cort =   se(cort, na.rm = TRUE)
  ) %>% gather("stat", "value", 1:length(.)) %>%
    mutate(value = round(value, 4)) %>%
    separate(stat, c("stat", "hormone")) %>%
    spread(stat, value)
```

```
## # A tibble: 2 x 4
##   hormone     mean      sd     se
## *   <chr>    <dbl>   <dbl>  <dbl>
## 1    cort   0.1877  0.1075 0.0066
## 2    test 177.5396 42.1505 2.6090
```

### Intrasexual Competitiveness Analyses

#### Descriptives

```
data_scaled %>%
  summarise(
    mean = mean(intr_cmpt, na.rm = TRUE),
    sd   = sd(intr_cmpt, na.rm = TRUE),
    se   = se(intr_cmpt, na.rm = TRUE)
  ) %>%
  mutate_all(round, 4)
```

```
## # A tibble: 1 x 3
##     mean     sd     se
##    <dbl>  <dbl>  <dbl>
## 1 2.9534 0.9847 0.0607
```

#### Cronbach's Alpha

```
horm_alphas <- data_hormones %>%
  select(ICS1:ICS12) %>%
  alpha()

horm_alphas$total$raw_alpha
```

```
## [1] 0.8634041
```

#### LMEM Analysis

```
model.IC.TbyC <- lmer(intr_cmpt ~ 1 + test.s * cort.s +
                        avg_test.s * avg_cort.s +
                        (test.s * cort.s || hm_id),
                      data = data_scaled, REML = FALSE)
summary(model.IC.TbyC)
```

```
## Linear mixed model fit by maximum likelihood t-tests use Satterthwaite
##   approximations to degrees of freedom [lmerMod]
## Formula:
## intr_cmpt ~ 1 + test.s * cort.s + avg_test.s * avg_cort.s + ((1 |
##     hm_id) + (0 + test.s | hm_id) + (0 + cort.s | hm_id) + (0 +
##     test.s:cort.s | hm_id))
##    Data: data_scaled
##
##      AIC      BIC   logLik deviance df.resid
##    459.9    502.5   -217.9    435.9      246
##
## Scaled residuals:
##      Min       1Q   Median       3Q      Max
## -2.74196 -0.51653 -0.01937  0.50550  2.56552
##
## Random effects:
##  Groups   Name          Variance  Std.Dev.
```

```
##  hm_id    (Intercept)   7.217e-01 8.495e-01
##  hm_id.1  test.s        3.064e-01 5.536e-01
##  hm_id.2  cort.s        3.949e-01 6.284e-01
##  hm_id.3  test.s:cort.s 1.533e-11 3.916e-06
##  Residual               1.468e-01 3.831e-01
## Number of obs: 258, groups:  hm_id, 59
##
## Fixed effects:
##                         Estimate Std. Error        df t value Pr(>|t|)
## (Intercept)              3.03493    0.12320  59.39000  24.634   <2e-16 ***
## test.s                   0.05333    0.22668  38.28000   0.235    0.815
## cort.s                   0.09377    0.21610  34.62000   0.434    0.667
## avg_test.s               0.38921    0.64713  60.29000   0.601    0.550
## avg_cort.s              -0.22143    0.86546  61.43000  -0.256    0.799
## test.s:cort.s            2.56471    1.56636 162.41000   1.637    0.103
## avg_test.s:avg_cort.s   -4.42082    3.02851  59.54000  -1.460    0.150
## ---
## Signif. codes:  0 '***' 0.001 '**' 0.01 '*' 0.05 '.' 0.1 ' ' 1
##
## Correlation of Fixed Effects:
##             (Intr) test.s cort.s avg_t. avg_c. tst.:.
## test.s      -0.004
## cort.s       0.025 -0.209
## avg_test.s   0.054 -0.003  0.002
## avg_cort.s   0.085 -0.002  0.016 -0.443
## tst.s:crt.s -0.069  0.017 -0.198 -0.013 -0.029
## avg_tst.:_. -0.380  0.002 -0.012 -0.142 -0.211  0.021
```

### State Anxiety Analyses

#### Descriptives

```
data_scaled %>%
  group_by() %>%
  summarise(
      mean = mean(st_anx, na.rm = TRUE),
      sd   = sd(st_anx, na.rm = TRUE),
      se   = se(st_anx, na.rm = TRUE)
  ) %>%
  mutate_all(round, 4)
```

```
## # A tibble: 1 x 3
##      mean     sd    se
##     <dbl>  <dbl> <dbl>
## 1 36.1217 8.8062 0.543
```

#### LMEM Analysis

```
model.SA.TbyC <- lmer(st_anx ~ 1 + test.s * cort.s +
                        avg_test.s * avg_cort.s +
                        (test.s * cort.s || hm_id),
```

```r
                    data = data_scaled, REML = FALSE)
summary(model.SA.TbyC)
```

```
## Linear mixed model fit by maximum likelihood t-tests use Satterthwaite
##   approximations to degrees of freedom [lmerMod]
## Formula: st_anx ~ 1 + test.s * cort.s + avg_test.s * avg_cort.s + ((1 |
##     hm_id) + (0 + test.s | hm_id) + (0 + cort.s | hm_id) + (0 +
##     test.s:cort.s | hm_id))
##    Data: data_scaled
##
##      AIC      BIC   logLik deviance df.resid
##   1797.6   1840.2   -886.8   1773.6      246
##
## Scaled residuals:
##      Min       1Q   Median       3Q      Max
## -2.73890 -0.54520 -0.08963  0.40247  2.98275
##
## Random effects:
##  Groups   Name          Variance Std.Dev.
##  hm_id    (Intercept)     32.25   5.679
##  hm_id.1  test.s          91.40   9.560
##  hm_id.2  cort.s           0.00   0.000
##  hm_id.3  test.s:cort.s 1237.24  35.174
##  Residual                 38.55   6.209
## Number of obs: 258, groups:  hm_id, 59
##
## Fixed effects:
##                        Estimate Std. Error       df t value Pr(>|t|)
## (Intercept)             36.5557     0.9238  55.5000  39.569   <2e-16 ***
## test.s                   2.9672     3.6544  45.5900   0.812    0.421
## cort.s                   7.0999     2.8540 146.3300   2.488    0.014 *
## avg_test.s               5.6698     4.9084  57.7900   1.155    0.253
## avg_cort.s              -6.1653     6.6422  60.2500  -0.928    0.357
## test.s:cort.s            1.0943    24.5607  18.1800   0.045    0.965
## avg_test.s:avg_cort.s -26.5258    22.8242  56.5500  -1.162    0.250
## ---
## Signif. codes:  0 '***' 0.001 '**' 0.01 '*' 0.05 '.' 0.1 ' ' 1
##
## Correlation of Fixed Effects:
##             (Intr) test.s cort.s avg_t. avg_c. tst.:.
## test.s       0.002
## cort.s       0.051 -0.220
## avg_test.s   0.061  0.001 -0.002
## avg_cort.s   0.094 -0.005  0.033 -0.446
## tst.s:crt.s -0.127 -0.005 -0.229 -0.015 -0.060
## avg_tst.:_. -0.382  0.004 -0.028 -0.167 -0.204  0.042
```
